# ANALYSIS AND IMPLICATIONS OF COMPLIANCE IN JOINT BIOMECHANICS SUPERPOSITION TESTING

**DOI:** 10.1101/2024.12.10.627572

**Authors:** Callan M. Gillespie, Nicholas J. Haas, Tara F. Nagle, Robb W. Colbrunn

**Author notes:** 2111 E 96^th^ Street, Cleveland OH 44106. 2111 E 96th Street, Cleveland OH 44106.

## Abstract

To quantify the contribution of specific ligaments to overall joint movement, the principle of superposition has been used for nearly 30 years. This principle relies on using a robotic test system to move a biological joint to the same position before and after transecting a ligament. The difference in joint forces before and after transecting the ligament is assumed to be the transected ligament’s tension. However, the robotic test system’s ability to accurately return the joint to the commanded pose is dependent on the compliance of the system’s various components, which is often neglected. Accordingly, there were three objectives in this manuscript: (1) **Explain** the influence of system compliance on positioning error in superposition testing with a mathematical model, (2) **Quantify** the compliance of components within the test system and (3) **Provide a framework** to evaluate uncertainty in published superposition based in situ force measurements, and demonstrate it on published Anterior Cruciate Ligament (ACL) forces. A system stiffness model was derived to explain that compliance of test system components will cause the superposition method to underestimate ligament tension and stiffness. Based on typical test system component and joint stiffness ranges measured in this study, it was determined that with decreasing robot and/or bone stiffness, or increasing joint stiffness values, ligament load error could increase to values greater than 50%. Results indicate that experimentalists should (1) increase test system component stiffness relative to joint stiffness and/or (2) compensate for compliance induced deflection of the test system components.

## 1. INTRODUCTION

The use of robots to study the biomechanics of human cadaveric joints began in the early 1990s [1]. Just a few years after the early adoption of robots, the principle of superposition began to be applied to *in vitro* orthopedic biomechanical testing to quantify the contribution of ligaments to joint mechanics [2, 3]. In 1995, superposition testing was performed on a modified uniaxial testing machine with high fixture stiffness [2, 3]. However, by the following year superposition testing began to be performed by 6-degree-of-freedom (DOF) serial arm robots, with inherently lower fixture stiffness [4, 5]. The scope of these studies also expanded over the next several decades to include the use of the displacement measurements from these serial arm robots. The load-displacement relationship of various ligaments was examined in more detail and parameters such as *in-situ* ligament slack length and stiffness were also quantified [6-9]. Many of the ligament parameters elucidated by superposition testing are used by clinicians and medical device companies to validate models, inform clinical understanding, and impact the design of new medical devices for joint replacements, reconstructions or repairs [3, 6, 9-12]. For example, Kent et al. used superposition testing to better understand the load-displacement relationship of the anterior cruciate ligament (ACL) and other stabilizing passive structures in the knee with the goal of reducing the risk of ACL graft failure [8].

Conceptually, superposition testing is simple, but how it is implemented determines how accurately one can quantify ligament parameters of interest. That implementation is often in 6-DOF and subject to positioning error in both translation and rotation. Superposition testing is performed by applying a known overall load vector to a biological joint 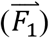 and recording the displacement vector of the joint relative to its neutral or zero load position (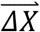, as illustrated in 1-DOF by the simplified spring model in **Figure 1**). After recording the displacement vector of the intact joint under the applied load, a structure within the specimen is then removed (e.g., Structure A in **Figure 1**), the joint is positioned back to the recorded displacement vector, and the new load is recorded 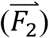. The new load 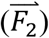 is the load carried by the remaining structures. The magnitude of the vector difference between the original and new load accounts for the load that was carried by the removed structure at that position 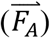 [2]. In the literature, all structures are generally ligaments whose removal involves transection of the ligament while loads are generally tension loads.

**Figure 1:**
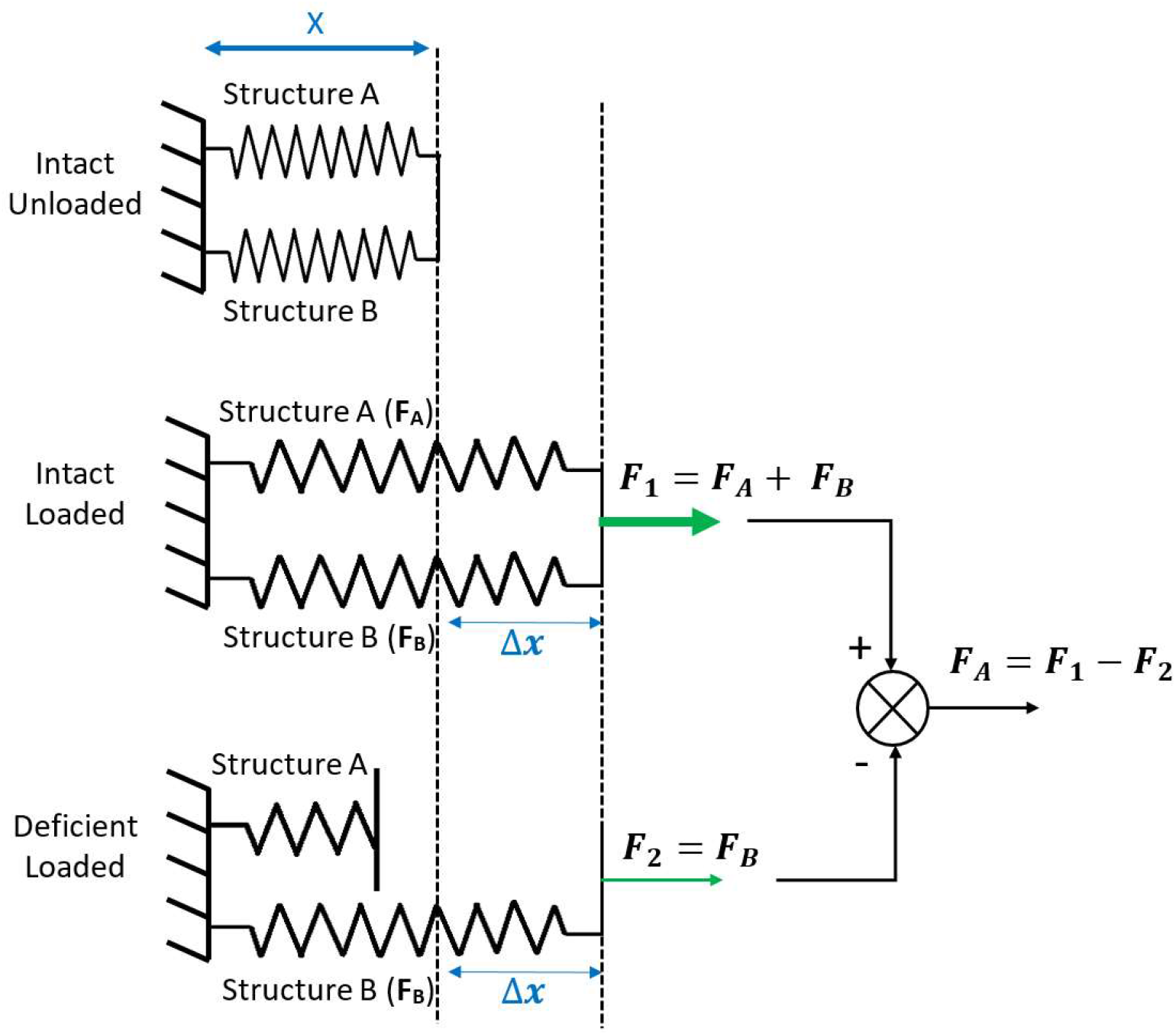
Simplified spring example of the Joint Stiffness Model used in traditional superposition testing. In this model, Structure A represents the structure whose contribution is being quantified using superposition, and Structure B represents all other structures in the joint.

Superposition testing is simplified to 1-DOF in the remainder of this paper for simplicity, but the underlying concepts also apply in 6-DOF. In the simplified spring model shown in **Figure 1**, Structure A represents the ligament whose contribution is being quantified, while Structure B represents all other ligaments and passive structures in the joint. These calculations follow the Joint Stiffness Model (JSM), which is traditionally used in superposition testing and assumes *ΔX* is identical between different ligament loading conditions.

This model assumes that *ΔX* is identical before and after Structure A (e.g., anterior cruciate ligament) is cut. However, errors in *ΔX* can be introduced if the compliance of the components in the test setup are not accounted for.

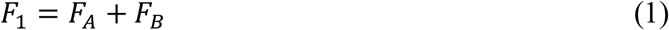

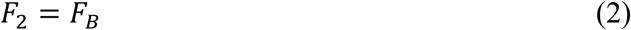

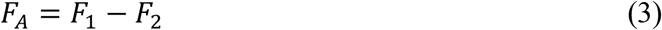

System displacement (*ΔX*), can only be identical between different surgical conditions if all the components of the test setup (except the biological joint) are infinitely rigid. Applying a load to the joint will result in loading every element in the test system (**Figure 2**). Components within the test system are not infinitely stiff, so with varying load observed in different surgical conditions, compliance in the test system will produce varying deflection, resulting in different *ΔX* values between surgical conditions.

**Figure 2:**
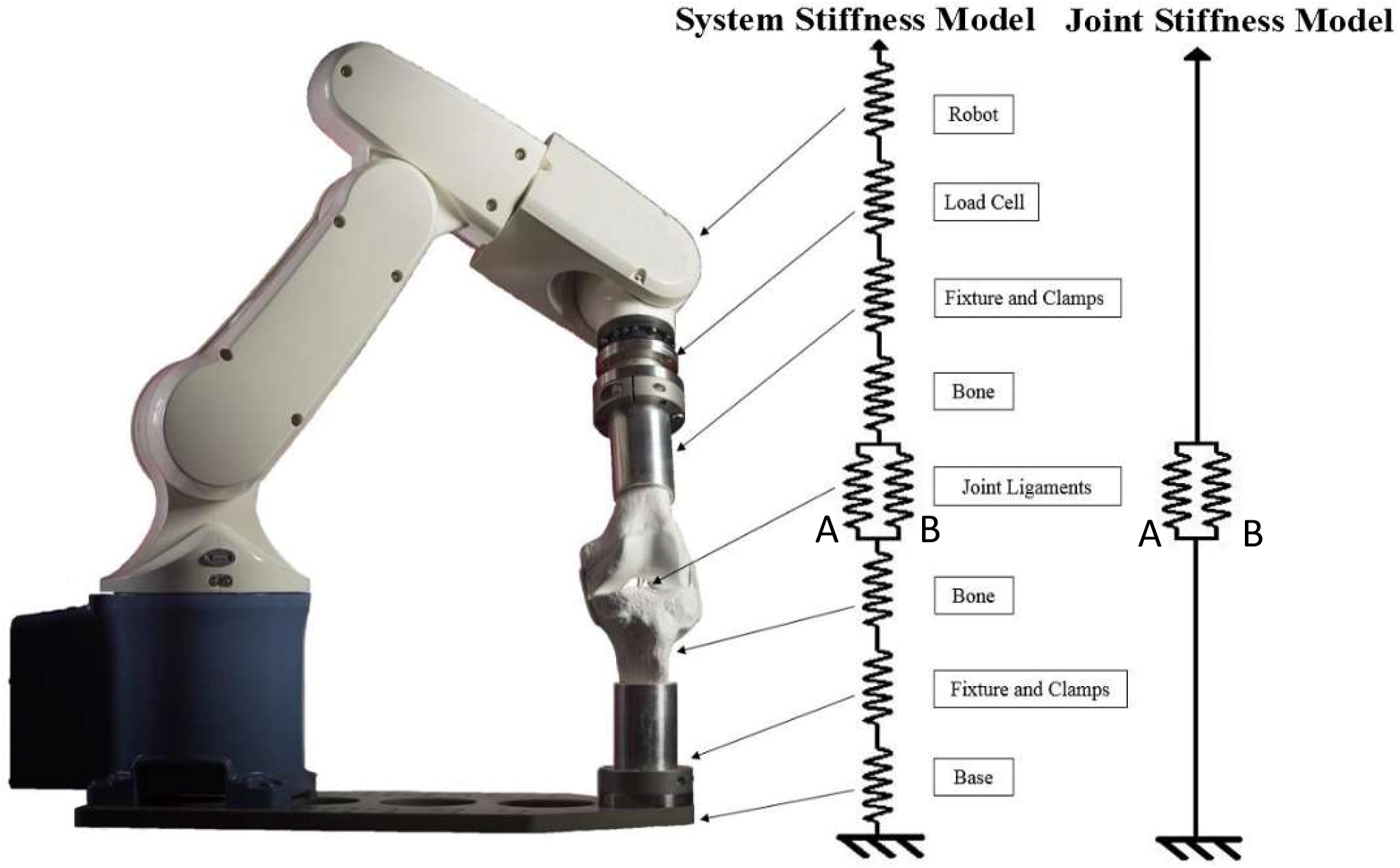
A typical robotic test system with the System Stiffness Model (SSM) and Joint Stiffness Model (JSM) superimposed. The SSM accounts for each component’s stiffness, while the JSM only accounts for the specimen’s stiffness.

While systemic errors in superposition have existed since its inception, it has been recognized that increased system compliance will result in increasing errors, and efforts have been made to minimize these errors. Fujie et al [13] developed a custom robotic system in 2004 that was designed to be stiffer than other 6-DOF robotic test systems and therefore reduce system deflection caused by compliance. Other groups have turned to parallel robots rather than serial arm robots in order to increase system stiffness at the cost of robot range of motion (ROM) limitations [14, 15].

To better understand the implications of the system stiffness assumptions within the JSM, a System Stiffness Model (SSM) is proposed, where each component of the system’s stiffness is included in the model, as seen in **Figure 2**. This includes the actuator (which can be a robot), load cell, fixtures, bone, and joint. The JSM equations (1-3) are rewritten with stiffness (*K*) and varying *ΔX* terms (4-6):

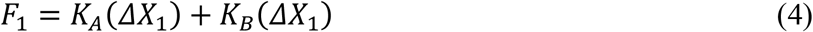

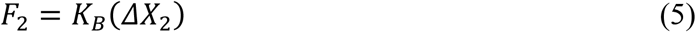

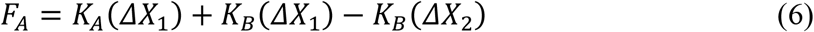

If *ΔX*_1_ ≠ *ΔX*_2_, then the resulting *F*_*B*_ terms, calculated as *K*_*B*_ (*ΔX*_1_) and *K*_*B*_ (*ΔX*_2_), respectively, do not cancel out to accurately calculate the load in Structure A. This will lead to systemic load error in the calculation of Structure A’s contribution to joint stability.

The mechanistic relationship between test system stiffness, positioning error, and the resulting systemic error in measuring both force and position within Structures A and B is explored further with a mathematical model of the SSM. By quantifying the stiffness of the test system components used in different laboratories, and applying those values into the SSM, the effect that positioning inaccuracy can have on resulting structure load contribution measurements in superposition testing can be estimated for different test systems. To this end, stiffness of each component (as seen in **Figure 2**) of a typical system must be quantified.

The objectives of this work were to: (1) Explain the mechanistic influence of system compliance on systemic positioning error in superposition testing with a mathematical model, (2) Quantify the stiffness of various components within different test systems and (3) Provide a framework by which to evaluate uncertainty in published in situ force measurements based on superposition testing, and demonstrate it on published ACL forces.

## 2. METHODS

### 2.1 System Stiffness Model (SSM)

To quantify the mechanistic influence system compliance has on positioning error, equations based on a simplified 1-DOF spring system were derived in detail in in Appendix 1. These equations, combined with the simplified spring model itself, make up the System Stiffness Model. The simplified 1-DOF spring system includes a Fixture Base (FB), Fixture Tool (FT) and Joint (J). The Fixture Base is rigidly mounted to the ground and includes components such as a test pedestal, mounting fixtures, bone, etc. The Fixture Tool manipulates the Joint and included components such as the robot, actuators, mounting fixtures, bone, etc. Similar to the JSM, the joint is made up of Structure A and B. Structure A represents the structure whose contribution is being quantified using superposition testing, and Structure B represents all other structures in the joint. **Figure 3** showcases this grouping and examines the effect compliance has on system displacement.

**Figure 3:**
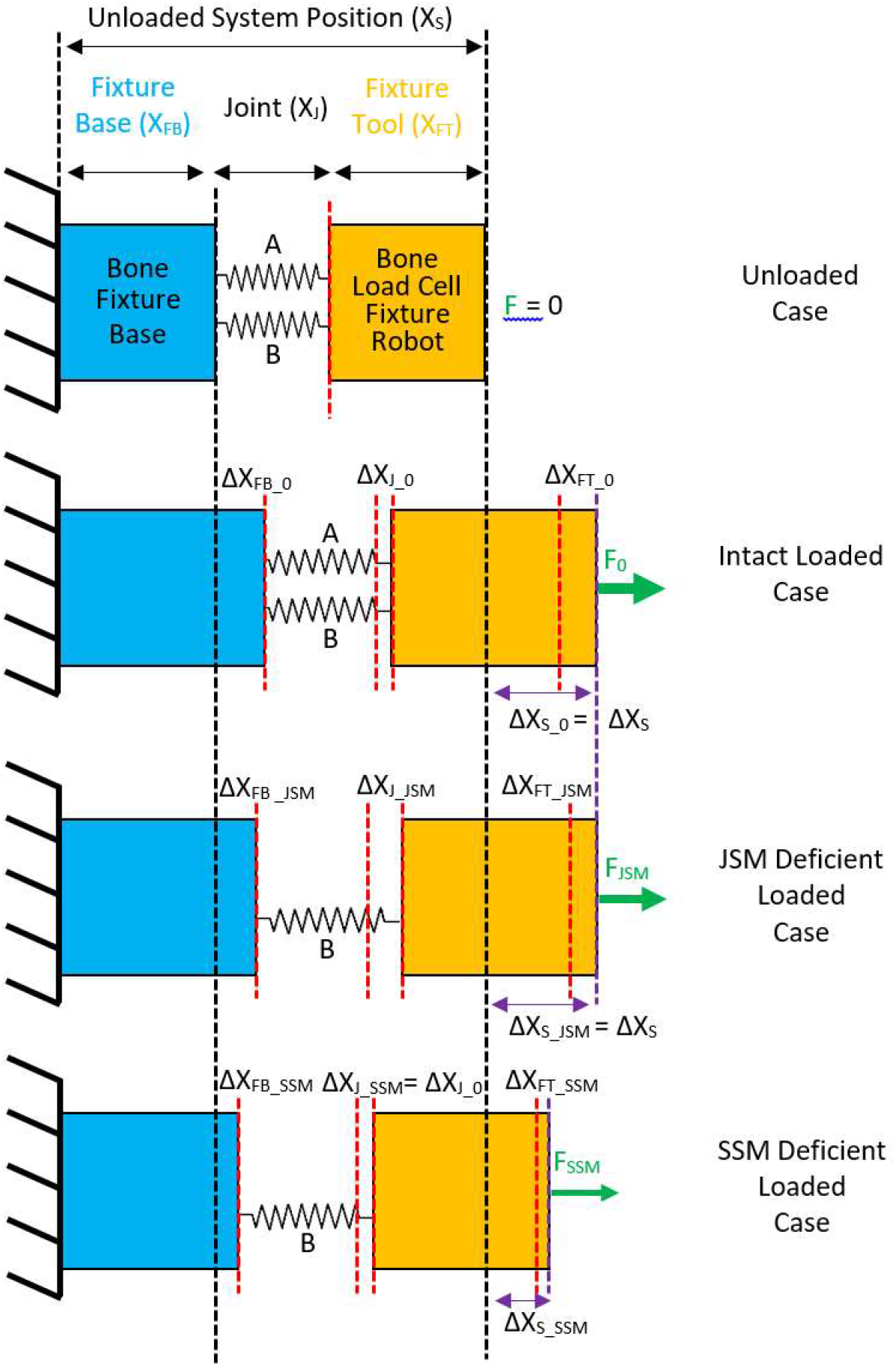
System Stiffness Model. The Intact System is positioned from an **Unloaded Case** to an **Intact Loaded Case** where initial system displacement is *ΔX*_*S*_0_ and initial joint displacement is *ΔX*_*J*_0_. The **JSM Deficient Loaded Case** demonstrates the distribution of system deflection due to the compliance of the various system components when the system is positioned to the same system displacement (*ΔX*_*S*_0_ = *ΔX*_*S*_*JSM*_). The **SSM Deficient Loaded Case** demonstrates the distribution of system deflection when the joint displacement is directly controlled (*ΔX*_*J*_), and the joint is brought to the same joint displacement*ΔX*_*J*_0_ = *ΔX*_*J*_*SSM*_.

*ΔX*_*S*_ is defined as the displacement of the system from its unloaded position (*X*_*S*_). The displacement of the Bone and Fixture Tool from their unloaded positions due to compliance are defined as *ΔX*_*FB*_ and *ΔX*_*FT*_, respectively. *ΔX*_*FB*_ and *ΔX*_*FT*_ are nonzero if there is deformation of the Fixture Base and Fixture Tool due to compliance. System displacement can be described as the sum of the displacements of all elements of the system, as given by (7):

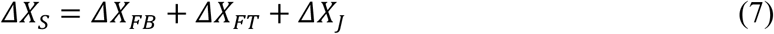

The JSM assumes that reported system displacement (*ΔX*_*S*_) is equal to joint displacement (*ΔX*_*J*_) because the Fixture Base and Fixture Tool deformation under applied load is neglected (i.e. *ΔX*_*FB*_ and *ΔX*_*FT*_ = 0). In the SSM, the Fixture Base and Fixture Tool deformation under applied load is included in the model, such that displacement of the system is modeled as springs in series that sum to *ΔX*_*S*_.

**Figure 3-Intact Loaded Case** showcases how in both the SSM and JSM, a known overall load is applied to the intact system (*F*_*O*_) and the testing system displacement (*ΔX*_*S*_0_) is measured (typically based on reported robot/actuator displacement). The joint displacement (*ΔX*_*J*_0_) can also be measured (typically based on displacement reported from Motion Capture). **Figure 3-JSM Deficient Loaded Case** showcases the JSM deficient loading state, where Structure A is cut and the system is commanded to go back to the same system displacement as in the Intact Loaded Case, (*ΔX*_*S*_0_ = *ΔX*_*S*_*JSM*_). Structure B is now the only structure resisting actuator motion. **Figure 3-SSM Deficient Loaded Case** showcases control of *ΔX*_*J*_ rather than *ΔX*_*S*_. Here *ΔX*_*J*_*SSM*_ is controlled to be the same as *ΔX*_*J*_0_, independent of fixture displacement.

Using the JSM and SSM model, equations were derived to understand whether published structure load contributions and stiffness values have been routinely over- or underestimated. While the SSM model makes predictions about both Structure A and B, superposition testing is generally used only to determine the properties of the ligament that is cut (i.e. Structure A). Therefore, the influence system stiffness has on load measurement error will focus on Structure A.

### 2.2 Quantification of System Stiffness

To quantify system stiffness, stiffness of each element in the system (as seen in **Figure 2**) was quantified. This included the load cell, custom fixtures, robot, bone, and joint ligaments. The human knee was chosen as the biological joint to quantify because it has been studied extensively with superposition testing in the literature [2-4, 6, 8, 10-13, 16, 17]. Therefore, the bones studied were the tibia and femur. The ACL was used as Structure A, and the remaining passive structures in the knee collectively made up Structure B. Stiffness measurements of the tibia and femur were constrained to the anterior-posterior (AP) direction because the ACL primarily resists anterior motion.

#### 2.2.1 Load Cell & Custom Fixture Stiffness

A representative load cell stiffness of 77,000 N/mm was obtained based on specifications for the ATI Omega 85 load cell (ATI, Apex NC). To allow the analysis to remain general purpose, lab specific custom fixtures were assumed to be infinitely rigid because of the range of variation from system to system. Many test systems have their own unique custom manufactured fixtures they use for testing and the effect of lab specific fixture stiffness should be calculated on a system by system basis.

#### 2.2.2 Robot Stiffness

Robot stiffness data was not easily accessible from the different manufacturers. As a result, robot stiffness was tested by the Cleveland Clinic BioRobotics Lab and collaborators at Kuka Systems North America, LLC. and University of Wisconsin-Madison. Seven different robots were tested to determine their respective stiffness. This was done by applying various static loads in the three base Cartesian axes of each robot using dead weights, ratchet straps and/or pulley systems while recording the displacement of the robot end-effector using an optical displacement measurement system. Increasing static loads were applied sequentially and care was taken to capture any stress relaxation of the applied load or creep of the robot end-effector. If there was creep and/or stress relaxation during loading then the robot displacement was averaged to account for this. While care was taken to ensure load was only along a single Cartesian axis of the robot, environmental factors sometimes made this difficult. Some robots were in rooms that prevented tension from being applied from a direction that was perfectly aligned with a robot’s Cartesian axis. In these cases, load was applied as close to the Cartesian axis as possible.

When loads were recorded by a 6-axis load cell attached to the robot end-effector, the resultant load was computed based on the 3 forces measured. Generally, robot displacements at low loads were beneath the noise floor of the optical displacement measurement sensors and, as a result, were ignored. The exception to this were lower payload robots, such as the Fanuc M-1iA Delta robot, which had measurable displacement even at lower loads. Applied loads were designed to cover a significant working range of the robot being tested. Multiple trials were conducted on the robots in order to obtain enough displacement and load values for reliable results.

Four different types of robots were tested: (1) parallel (Mikrolar robots), (2) serial arm (Kuka and Denso robots), (3) delta (Fanuc), and (4) a custom designed robot for joint biomechanics research [13]. Robots tested for use in this manuscript are outlined in Table 1 with experimental details. The data from the custom designed robot is based on previously published values from literature [13].

**Table 1:** Robot Stiffness Test Matrix.

| **Robot** | **Institute** | **Motion Capture System** | **Load Measurement** | **Resultant Loads Applied (N)** |
| --- | --- | --- | --- | --- |
| Mikrolar R2000 | Cleveland Clinic BioRobotics Lab | Optotrak Certus (NDI, Waterloo, ON) | ATI Theta | 200-900 |
| Mikrolar R2000 w Custom Rotary Stage | Cleveland Clinic BioRobotics Lab | Optotrak Certus | ATI Theta | 200-900 |
| Kuka KR300 Ultra 2500 | Kuka Systems North America | Laser Micrometer | Electronic Spring Scale | 500-1,500 |
| Kuka KR300 Ultra 2700-2 | University of Wisconsin-Madison | Optitrack w Prime<sup>x</sup> 13 (Corvalis Oregon) | ATI Omega 160 | 20-800 |
| Custom Robot [13] | Kogakuin University | Laser Transducer | UFS-6520A250 | 0-500 |
|  |  | (Keyence, Osaka Japan) | (Osaka, Japan) |  |
| Denso VP-6242G** | Cleveland Clinic BioRobotics Lab | Optotrak Certus | ATI Mini 45 | 5-60 |
| Kuka KR16 | Cleveland Clinic BioRobotics Lab | Optotrak Certus | ATI Delta | 40-800 |
| Fanuc M-1iA | Cleveland Clinic BioRobotics Lab | Optotrak Certus | Dead weights with Pulley | 0-22 |

Robot stiffness is a function of both loading direction and robot pose, so care was taken to be as consistent as possible. Each robot was brought to a respective standard pose to better compare results. All poses for serial arm robots were similar, where all links were oriented vertically or horizontally, with the general pose in a “hang-man” orientation. The parallel robots’ home position was defined by a position where the top plate’s was parallel to its base, and the plate’s height was positioned at the middle of its range. The stiffness was calculated in each Cartesian direction of the robot’s base frame by dividing the resultant force applied by the resultant measured displacement. During testing the robot drives were turned on so that robot stiffness was similar to an actual test rather than a function of a robot’s brakes.

#### 2.2.3 Bone Stiffness

The femur and tibia stiffness were computed by measuring displacement of the bones (*ΔX*_*Bone*_) using an electronic test indicator (Fowler, Boston MA) while applying a cantilever load (*P*) to the bone as close to the joint as possible with a uniaxial testing system (MTS 858, Eden Prairie MN) as seen in **Figure 4**. One tibia with an approximate length (*l*) of 12.4 cm and one femur with a length of 15.1 cm were loaded in the anterior and posterior directions ranging from 50 to 300 N in increments of 50 N. The load (*P*) was plotted as a function of displacement (*ΔX*_*Bone*_), and the slope of the linear regression equation was used as the stiffness value in the anterior and posterior directions. The flexural rigidity (*EI*) was calculated based on the equation for a cantilever beam’s maximum deflection with a load at the end (8). The flexural rigidity of each load and AP orientation combination was then averaged for each bone to calculate a mean value for flexural rigidity per bone.

**Figure 4:**
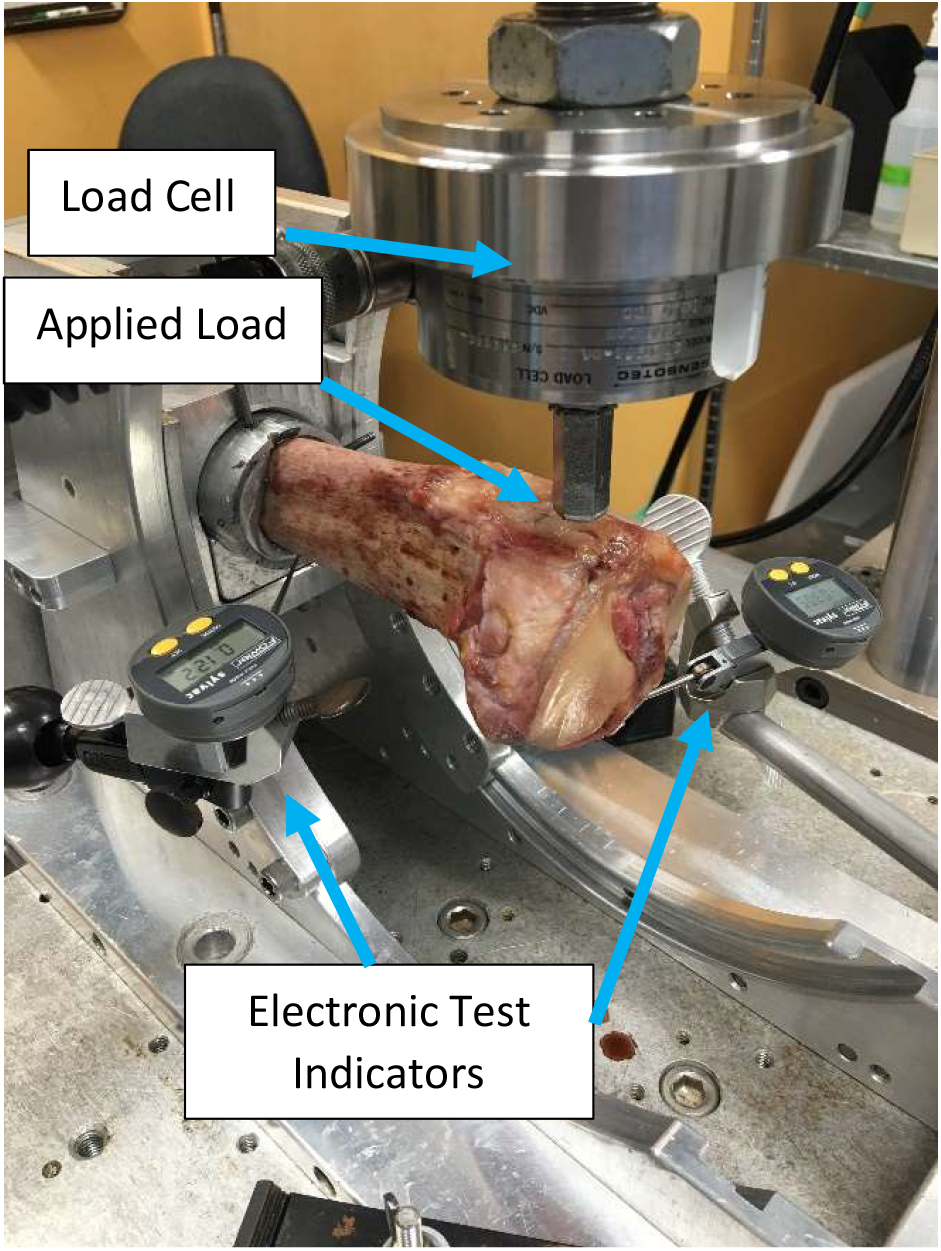
Bone stiffness measurement setup, including uniaxial machine for applying loads to the bones in the knee in the anterior and posterior directions (photo displays tibia) and electronic test indicators for measuring bone deflection.

The calculated values for flexural rigidity were compared to values from literature for validation [18]. Cantilever stiffness can be rewritten in terms of flexural rigidity of the given bone and the length of that bone extending out of the fixture (9). Therefore, bone stiffness values in the AP direction were also computed based on literature and compared to experimentally measured values. It should be noted that in (9) as bone length increases, AP stiffness decays exponentially.

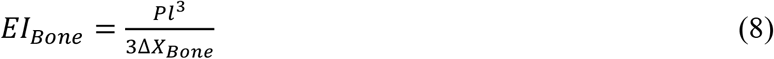

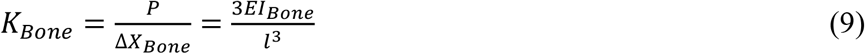

#### 2.2.4 Joint Stiffness

The knee joint AP stiffness was calculated using data from the <u>Open Knee project</u> [19]. Joint kinematic data was calculated from reported robot pose, but the influence of compliance was minimized by using a very stiff robot (the Mikrolar R2000 robot with stage). The AP laxity data from seven specimens was collected at 0, 30, 60 and 90 degrees of flexion with applied anterior forces ranging from -100 to 100 N in increments of 10 N. At each flexion angle the data from the seven knees was averaged and AP displacement from neutral was plotted against anterior force (**Figure 6A**). Unlike robot and bone stiffness, joint stiffness was not considered to be linear. Third order polynomial functions were fit to the average laxity data at each flexion angle. By taking the first derivative of these polynomials, AP stiffness was calculated. Stiffness was plotted against anterior displacement and force in order to directly compare joint stiffness to the stiffness of other components in test systems.

### 2.3 Computing Load Measurement Error

To estimate the effect that positioning error can have on Structure A (i.e. ACL) load measurement error in a knee model, it was necessary to (1) compute positioning error as a function of system component stiffness to establish a newly defined compliance compensation (*CC*) metric, and (2) use the Compliance Compensation Metric to calculate System Load Error.

#### 2.3.1 Compliance Compensation Metric

The positioning error when superposition testing is performed without correcting for compliance is predicted by the SSM and is depicted in **Figure 3** Intact Loaded and JSM Deficient Loaded cases. In both cases, the system displacement does not equal the joint displacement (*ΔX*_*S*_ ≠ *ΔX*_*J*_) due to compliance. More importantly, the joint displacement between the two cases is also not equal (*ΔX*_*J*_0_ ≠ *ΔX*_*J*_*JSM*_). If the joint displacements were equal then the principle of superposition would be satisfied. The Compliance Compensation Metric aims to quantify the joint positioning error predicted by the SSM when system compliance is not accounted for. In other words, the Compliance Compensation Metric is the difference in joint displacement between the Intact Loaded and JSM Deficient Loaded cases:

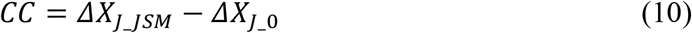

To calculate the Compliance Compensation Metric, *ΔX*_*J*_*JSM*_ and *ΔX*_*J*_0_ need to be solved for as a function of component stiffness. Rewriting (7):

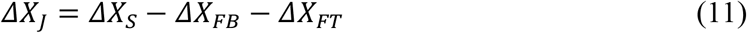

Because the tibia (*Tib*) and load cell (*LC*) are often connected in series to the ground, while the femur (*Fem*) and robot (*Rob*) are connected in series to make up the tool in **Figure 3**; *ΔX*_*FB*_ and *ΔX*_*FT*_ can be rewritten as:

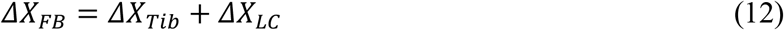

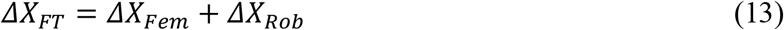

Therefore joint displacement is system displacement minus each component’s compliance induced displacement:

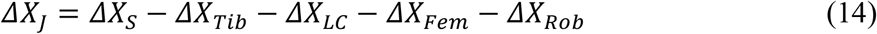

In the SSM, each component is modeled as a spring, and the linear spring equation (*ΔX* = *F*/*K*) may be substituted into (14):

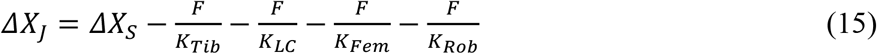

Substituting (15) into Compliance Compensation Metric (10), gives

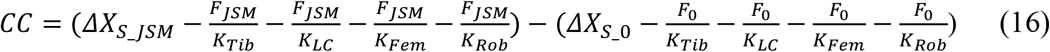

In (16) the *ΔX*_*S*_ terms cancel, and it can be simplified further:

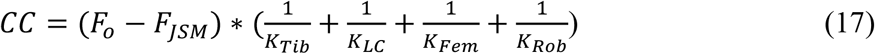

Finally, by substituting (9) into (17) for the femur and tibia stiffness, bone stiffness can be written in terms of bone length:

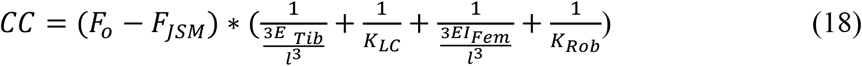

The Compliance Compensation Metric in (18) can therefore be written solely as a function of applied load (*F*) and component system stiffness or bone length (*l*). By varying robot stiffness and bone length, the effect of component stiffness on positioning error can be better understood.

In (18) it should be noted that only the flexural rigidity (*EI*) values from literature (**Table 3**) were used. The tibia and femur shared the same bone length (*l*) because it was assumed that the tibia and femur were potted at the same distance from the joint line. However, a range of bone lengths (*l*) were used between 5 and 30 cm that reflected typical bone testing lengths. Similarly, a range of robot stiffness values were used between 50 and 2000 N/mm that reflected typical robot stiffness (**Table 2**). A previously published load cell stiffness value for an ATI Omega 85 Load Cell was also used as a representation for all load cells.

**Table 2:** Robot Stiffness.

| Robot | Stiffness (N/mm) |  |  |  |
| --- | --- | --- | --- | --- |
|  | X axis | Y axis | Z axis | Average |
| Mikrolar R2000 | 2883 | 4733 | 16622 | **8079** |
| Mikrolar R2000 w Stage [14] | 1148 | 2198 | 9523 | **4290** |
| Kuka KR300 Ultra 2500 | 1707 | 731 | 1286 | **1241** |
| Kuka KR300 Ultra 2700-2 | 1223 | 557 | 701 | **827** |
| Custom Robot [13] | 333 | 1000 | 1000 | **778** |
| Denso VP-6242G** | 40 | 520 | 980 | **512** |
| Kuka KR16 | 181 | 249 | 237 | **222** |
| Fanuc M-1iA | 59 | 21 | 10 | **30** |

**Table 3:**
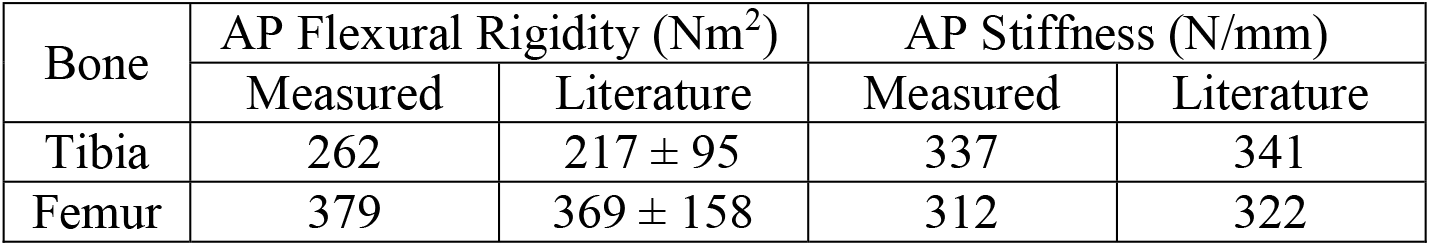
Flexural rigidity and bone stiffness data.

| Bone | AP Flexural Rigidity (Nm<sup>2</sup>) |  | AP Stiffness (N/mm) |  |
| --- | --- | --- | --- | --- |
|  | Measured | Literature | Measured | Literature |
| Tibia | 262 | 217 ± 95 | 337 | 341 |
| Femur | 379 | 369 ± 158 | 312 | 322 |

#### 2.3.2 System Load Error Calculation

To estimate the effect that positioning error can have on Structure A (i.e. ACL) load measurement uncertainty, it is first necessary to define load measurement error as function of positioning error. To do this, ACL tension (*F*_*A*_ from **Figure 1)** was calculated for the JSM vs SSM cases in **Figure 3**. Normalized System Load Error 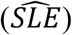 was defined as:

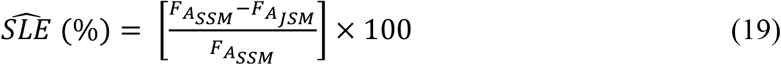

Values for ACL load (*F*_*A*_*JSM*_) contribution in the JSM case, as a function of initial load (*F*_0_) or system displacement (*ΔX*_*S*_), have been reported extensively in literature. However, the corresponding ACL load (*F*_*A*_*SSM*_) in the SSM case, is unknown. Therefore, it was necessary to develop a methodology to estimate what the ACL loads (*F*_*A*_*SSM*_) would be if system compliance is compensated for during superposition testing. To calculate *F*_*A*_*SSM*_ it is first necessary to rewrite (19) by substituting the ligament force terms similar to (3).

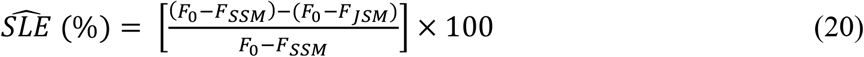

The only unknown term (20) was *F*_*SSM*_, the restraint force the Structure A deficient knee would have had if the joint were positioned to the same joint displacement between intact and Structure A deficient cases, and system compliance were compensated for (i.e. *ΔX*_*J*_0_ = *ΔX*_*J*_*SSM*_). The approximation of *F*_*SSM*_ involved four parts: (1) Obtaining the value of *F*_JSM_ from literature, (2) Solving for *ΔX*_*J*_*JSM*_ using the model of knee joint laxity, (3) Calculating the Compliance Compensation Metric, and (4) Solving for *ΔX*_*J*_*SSM*_ using the model for knee joint laxity. The methodology is shown graphically below (**Figure 5**).

**Figure 5:**
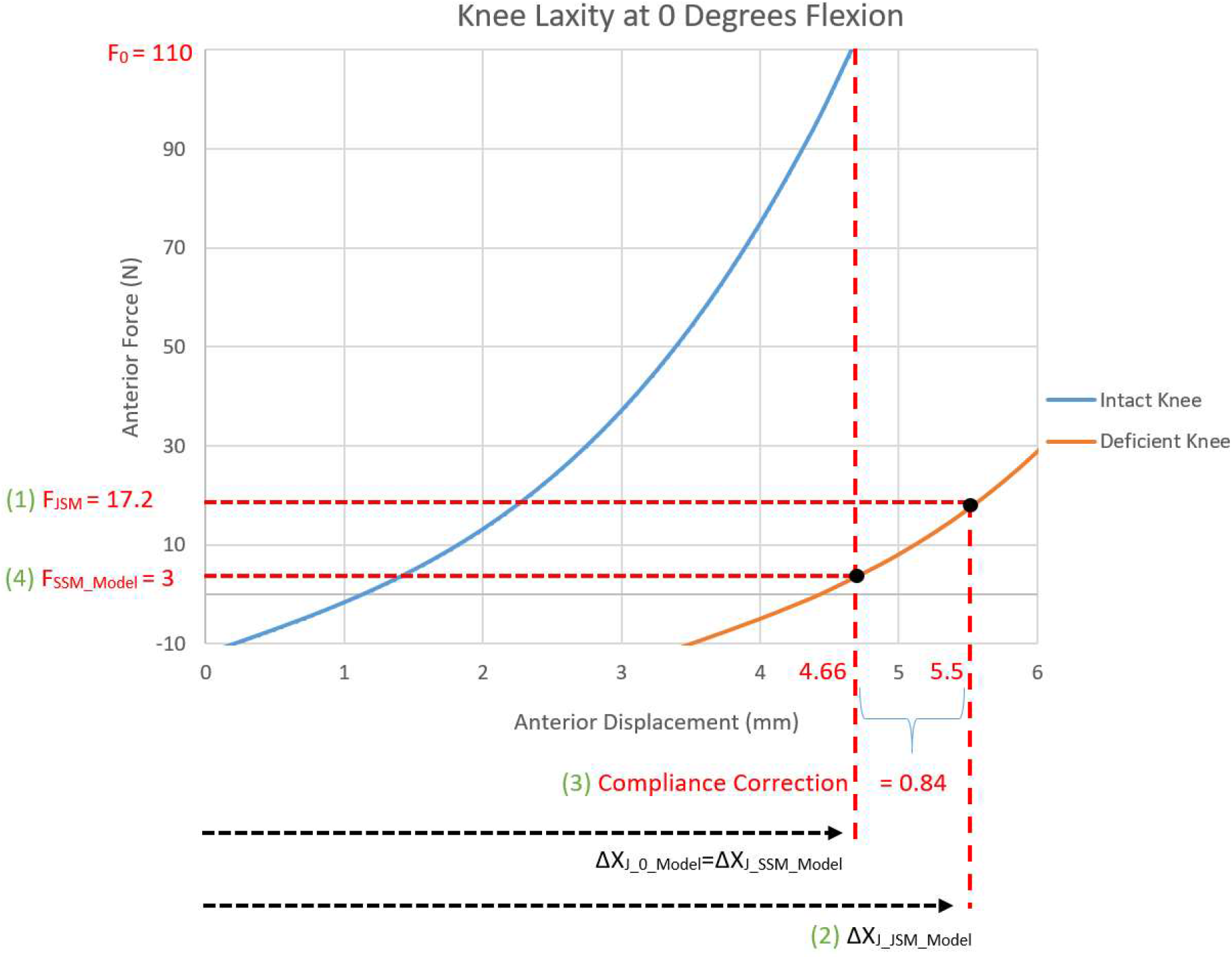
The methodology used to solve for *F*_*SSM*_. (1) The force applied to the ACL deficient knee (*F*_*JSM*_) was extracted from literature. (2) The joint displacement at the ACL deficient force (*ΔX*_*J_JSM_Model*_)was solved for using the model of knee laxity as described by a 3^rd^ order polynomial in Open Knees data [19]. (3) A Compliance Compensation Metric was applied to *ΔX*_*J_JSM_Model*_ in order to determine what joint deflection (*ΔX*_*J_*0*_Model*_ *= ΔX*_*J_SSM_Model*_) would have been at the initial load (*F*_0_). (4) At the corrected joint displacement, a corrected joint force (*F*_*SSM_Mod*_) was calculated. *F*_*SSM_Model*_ was calculated for a variety of knee flexion angles, robot stiffness, and bone lengths. This figure displays an example where the knee joint was positioned at 0 degrees of flexion, the robot stiffness was 1241 N/mm, (equivalent to a Kuka KR300 Ultra 2500 robot), and the bone length was 15 cm.

To demonstrate the framework for evaluating uncertainty in published in situ force measurements, data from Sakane et al. was utilized, where 110 N of anterior drawer (*F*_0_), was applied to 10 intact knees, followed by superposition testing of the paired ACL deficient knees to compute the mean ACL Load (*F*_*A*_) at knee flexion angles of 0, 30, 60, and 90 degrees [2, 16]. By subtracting published values for *F*_*A*_ from *F*_0_, as in (3), *F*_*JSM*_ was approximated and used to calculate joint displacement from a nonlinear 3^rd^ order polynomial joint stiffness model of the knee joint, elucidated from the Open Knees project (**Figure 6**) [19]. Using the polynomial equation to solve for a cubic root, the joint displacement *ΔX*_*J_JSM_Model*_ was solved for at the reported ACL deficient force *F*_*JSM*_. Polynomial joint stiffness models were derived at each included flexion angle (0, 30, 60, and 90 degrees), because knee laxity differs as a function of flexion angle.

**Figure 6:**
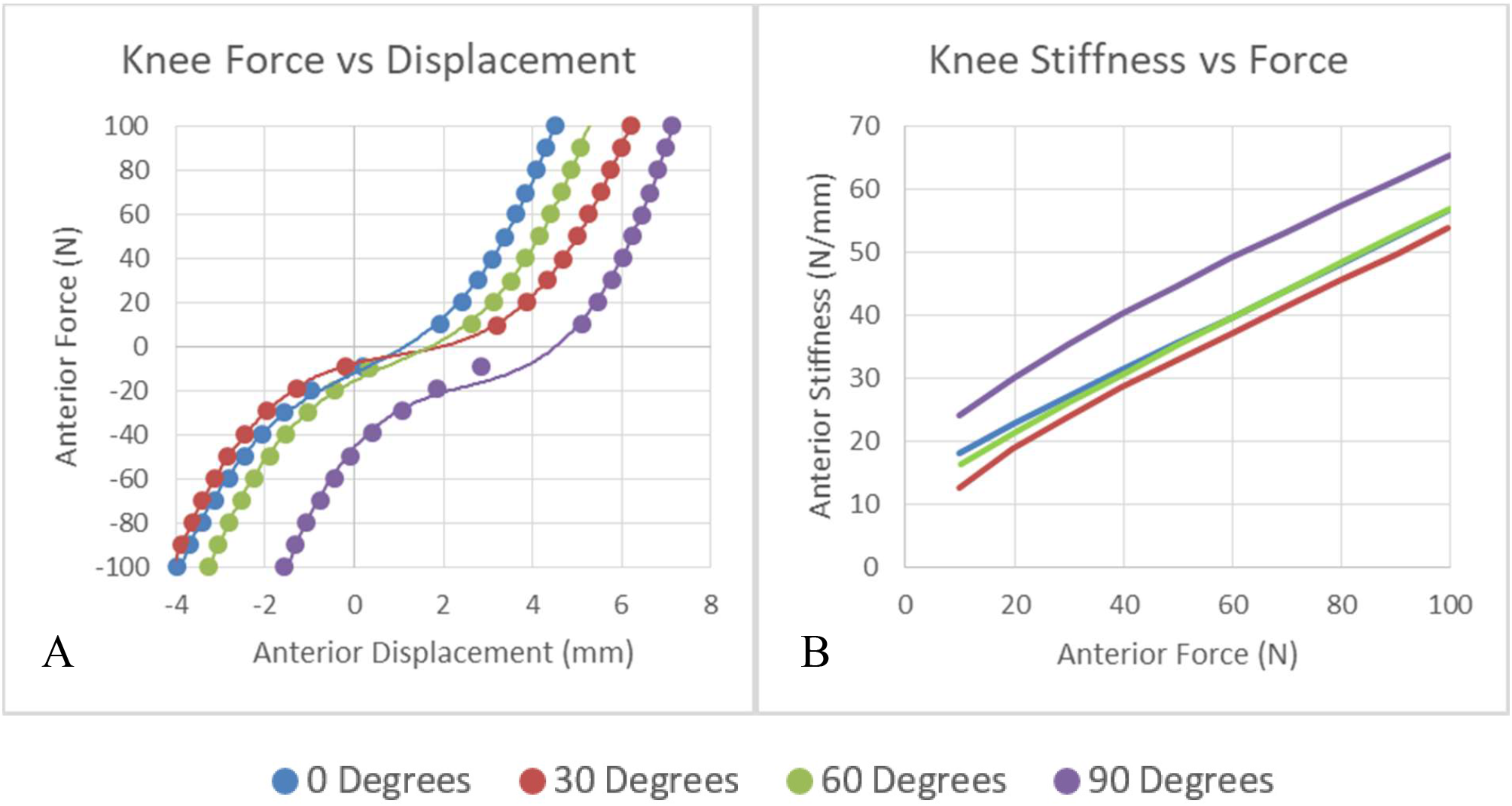
A) showcases the AP Force-Displacement relationship for the knee joint (Anterior Drawer Force and Anterior Translation are positive). B) Showcases how stiffness changes as a function of force. Both figures display results for all included flexion angles.

To compensate for the compliance induced positioning error in *ΔX*_*J_JSM_Model*_, the compliance correction term was used. The compliance correction term varied based on F_0_ and F_JSM_ from Sakane et al. and robot and bone stiffness (18) [16].

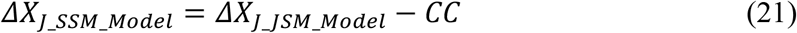

Once *ΔX*_*J_SSM_Model*_ was calculated, it was applied to the polynomial AP stiffness model for a specific flexion angle to calculate *F*_*SSM_Model*_ at that flexion angle, as shown in **Figure 5**. System Load Error at that flexion angle was then calculated by applying *F*_*SSM_Model*_ into (20) for *F*_*SSM*_. System Load Error in the knee model represented the % error in ACL tension load due to systemic positioning error in superposition testing. A range of bone lengths and robot stiffness data was used to calculate a range of System Load Errors on a 3D graph, with the x-axis representing robot stiffness, the y-axis representing bone stiffness, and the z-axis representing % System Load Error.

## 3. RESULTS

### 3.1 System Stiffness Model

In addition to providing an intuitive framework to understand system compliance, the SSM also provides insight into how system positioning errors can affect superposition computed ligament tensions when system compliance is not compensated for. In the Intact Loaded Case in **Figure 3** there are 2 ligaments resisting system displacement whereas in the JSM Deficient Loaded Case there is only one. The decrease in resistance in the joint to the same desired system displacement results in a decreased force *F*, where *F*_*JSM*_ < *F*_0_. Reduced applied force with identical fixture stiffness results in reduced fixture deformation after the structure is cut, and therefore, changes to the distribution of the system displacement. Fixture displacement, *ΔX*_*FB*_ and *ΔX*_*FT*_, will reduce, such that *ΔX*_*FB_JSM*_ < *ΔX*_*FB_*0_ and *ΔX*_*FT_JSM*_ < *ΔX*_*FT_*0_, which will result in increased joint displacement *ΔX*_*J*_, where *X*_*J_JSM*_ > *ΔX*_*J_*0_. A detailed derivation of this may be found in Appendix 1.

Going back to equations 1-6, but now focusing on joint displacement rather than system displacement, the *ΔX*_*J*_ term can be substituted for *ΔX*:

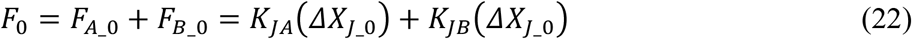

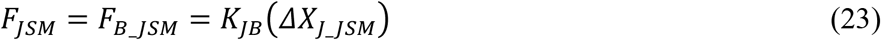

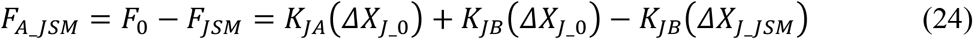

In (22-24) the *K*_*JA*_ and *K*_*JB*_ terms are written differently from the *K*_*A*_ and *K*_*B*_ terms in (4-6) because they have a different physical meaning. The *K*_*A*_ and *K*_*B*_ stiffness terms are derived from the JSM, and because that model doesn’t distinguish between system and joint stiffness, the JSM stiffness terms are a function of both the fixture and structure stiffness. The *K*_*JA*_ and *K*_*JB*_ terms are derived from the SSM and are purely a function of the stiffness of the A & B structures respectively. As in (6), the *F*_*B*_ terms, calculated as *K*_*JB*_ (*ΔX*_*J_*0_) and *K*_*JB*_(*ΔX*_*J_JSM*_), don’t cancel because *ΔX*_*J_*0_ ≠ *ΔX*_*J_JSM*_. Further, if *X*_*J_JSM*_ > *ΔX*_*J_*0_, then *F*_*JSM*_ in (23) is an overestimation for Structure B’s force contribution, and has been historically overestimated when system compliance was not compensated for. Examining (24), if *F*_*JSM*_ is overestimated then Structure A’s force, *F*_0_ − *F*_*JSM*_, will be underestimated (3) by the JSM.

The SSM can also provide insight into ligament stiffness parameter errors computed based off superposition computed tension and reported system displacement. This derivation can be found in Appendix 2, where it is derived that system compliance causes Structure B stiffness to be underestimated (A2-4). Further, (A2-12) indicates that Structure A stiffness is also consistently underestimated by the JSM method.

If joint displacement rather than system displacement were controlled, as in the SSM Deficient Loaded Case in **Figure 3**, then *ΔX*_*J_SSM*_ would replace *ΔX*_*J_JSM*_ in (24):

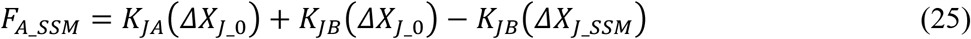

Because *ΔX*_*J_*0_ = *ΔX*_*J_SSM*_, the *K*_*JB*_ terms in (25) cancel out and reduce to (3). This satisfies the fundamental superposition assumption made by the JSM and brings the two models back into agreement.

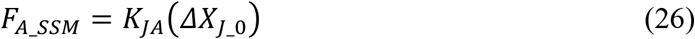

To better understand how the magnitude of error in estimating Structure A tension with the superposition method is related to system compliance, the absolute System Load Error (*SLE*), the numerator in (19), was interrogated in greater detail.

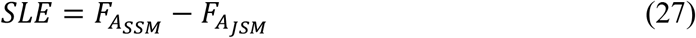

Substituting (26) and (24) gives the equation,

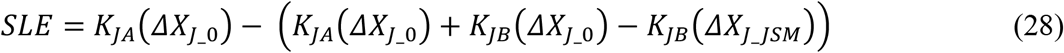

The K_JA_ terms cancel out and simplify.

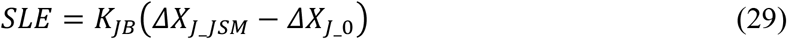

Equation (29) can be rewritten in terms of the Compliance Compensation Metric,

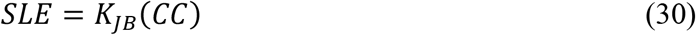

The Compliance Compensation Metric quantifies the joint positioning error predicted by the SSM when system compliance is not accounted for. Therefore equation (30) demonstrates that System Load Error is proportional to Structure B stiffness and the magnitude of the joint positioning error. The Compliance Compensation Metric can be rewritten given (17) and (A1-3), giving:

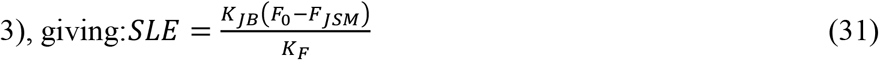

In (31), *K*_*F*_ is the resultant stiffness of all fixtures as given by (A1-3). Since *F*_0_ > *F*_*JSM*_, equation (31) demonstrates that System Load Error is also inversely proportional to fixture stiffness. Therefore the ratio of Structure B joint stiffness to fixture stiffness is proportional to the System Load Error.

### 3.2 Quantification of System Stiffness

The load cell, lab specific fixtures (such as clamps), robot, bone, and joint all make up system stiffness, as depicted in **Figure 2**.

#### 3.2.1 Robot Stiffness

Stiffness measurements from all robotic systems detailed in Table 1 are presented in Table 2.

There were large differences in robot stiffness depending on the direction of loading. However, in the three translational DOF, the relationship between applied load and robot displacement was found to be linear for each robot tested by the Cleveland Clinic BioRobotics Lab or collaborators.

#### 3.2.2 Bone Stiffness

Flexural rigidity and stiffness of the tibia and femur were calculated from loading conditions in the AP direction and compared with data from literature [18]. The results are presented in Table 3.

The AP stiffness values presented in **Table 3** correspond to the bone lengths measured by Cleveland Clinic BioRobotics Lab: a tibia of 12.4 cm and femur of 15.1 cm. Due to good agreement between literature and experimental data, the AP flexural rigidity values from literature were used to calculate System Load Error.

#### 3.2.3 Joint Stiffness

Average AP translation from the seven specimens was plotted against force for each included flexion angle, **Figure 6A**. Knee stiffness as a function of anterior displacement and force is also displayed in **Figure 6B**, to directly compare joint stiffness to other system stiffness.

### 3.3 System Load Error

*F*_*JSM*_ was calculated as 17.2, 19.1, 40, and 50.7 N for 0, 30, 60, and 90 degrees flexion, respectively. Error computed as a function of robot stiffness and bone length is shown in **Figure 7** for 0, 30, 60, and 90 degrees flexion, displaying that as robot stiffness decreases or as bone length increases, the error in ACL contribution increases.

**Figure 7:**
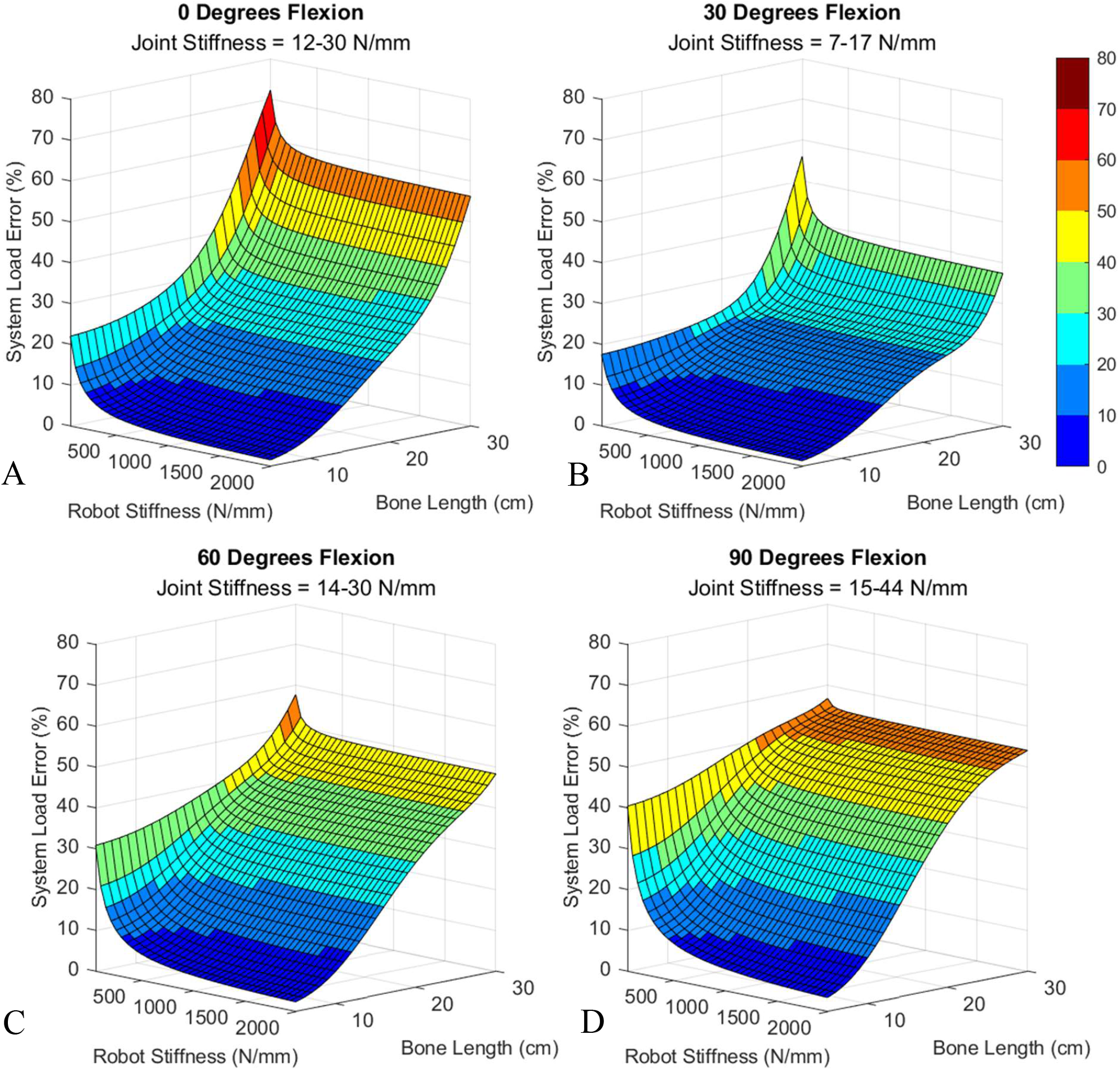
Estimated ACL Contribution Error as a function of Robot Stiffness and Bone Length. **A-D** showcase System Load Error for the different knee flexion angles that laxity tests were conducted at. The range of joint stiffness computed from the knee model during the System Load Error calculation is also shown.

For bone lengths ranging from 5 to 30 cm, and robot stiffness values ranging from 50 to 2000 N/mm, for flexion angles of 0, 30, 60 and 90 degrees, respectively, System Load Errors ranging from 1 to 72 percent were observed across all flexion angles included. The System Load Error at the highest joint stiffness range (90 degrees) was greater than at the lowest joint stiffness range (30 degrees) for all matched combinations of robot stiffness and bone length.

## 4. DISCUSSION

There were 5 key findings from this study. 1) Fixture compliance causes the principle of superposition to be violated. 2) When compliance is not taken into account, as in the JSM model (**Figure 1)**, Structure A load is underestimated, Structure B load is overestimated, and structure stiffness is underestimated. 3) The magnitude of system error is proportional to the ratio of Structure B joint stiffness to fixture stiffness and the amount of compliance as given by the compliance correction term. 4) The magnitude of superposition computed System Load Errors can be substantial with some values observed over 50%. 5) By directly controlling joint displacement (*ΔX*_*J*_) rather than system displacement (*ΔX*_*S*_) the principle of superposition is satisfied and the effect of fixture compliance on System Load Errors can be eliminated.

### 4.1 System Stiffness Model

Describing test system components as springs in series and in parallel presents a model for accounting for the error induced by system compliance when enforcing the fundamental assumption of superposition (**Figure 2**). The SSM shows that *ΔX*_*S*_ ≠ *ΔX*_*J*_ when there is system compliance. The magnitude of that disagreement is given by the Compliance Compensation Metric (10), which quantifies the joint positioning error predicted by the SSM when system compliance is not accounted. In addition, the SSM predicts that historically there has been:

1. Underestimation of Structure A tension
2. Overestimation of Structure B tension
3. Underestimation of both Structure A & B stiffness values (Appendix 2)

The System Load Error in Structure A was analyzed in more detail because it was assumed to be the primary structure of interest in most superposition studies. Equations (30-31) describe Structure A System Load Error and demonstrate that, in addition to the Compliance Compensation Metric, fixture stiffness (such as robot and bone) and joint stiffness also influence overall error. From equation (31), increasing joint stiffness (*K*_*JB*_) or reducing fixture stiffness (*K*_*F*_) will result in increasing System Load Error in superposition testing. If System Load Error is to be reduced then the ratio of joint stiffness to fixture stiffness should be minimized. Another option to reduce systemic error is to directly control joint displacement (*ΔX*_*J*_), which would ultimately reduce the Compliance Compensation Metric in (30) to negligible values. Directly controlling joint displacement is functionally equivalent to having infinite fixture stiffness, which is borne out in equations (30) and (31). If the Compliance Correction Term in (30) is minimized it implies that the effective fixture stiffness in (31) goes towards infinity.

The effect fixture stiffness/bone length versus joint stiffness has on System Load Error can be understood from **Figure 7**, where the individual plots indicate increasing error with decreasing fixture stiffness and increasing bone length and the overall error levels are greater in plots with higher joint stiffness flexion angles (i.e. 90 degrees) compared to plots with lower joint stiffness flexion angles (i.e. 30 degrees). This provides evidence for why robot stiffness has been prioritized in past superposition testing [13].

It should be noted that while the SSM does simplify a test system into only elastic elements in order to better understand the effect of system compliance on systemic load and stiffness error. Additional sources of error due to viscoelastic effects, control algorithm setpoint tracking, friction, and backlash are not taken into account in the SSM.

### 4.2 System Stiffness

#### 4.2.1 Robot Stiffness

In this study, the stiffness of eight different robots were obtained and stiffness values varied greatly. While selecting the highest stiffness robot might seem like the best solution to reducing System Load Error in superposition testing, there are other attributes that should be considered. The robot must have sufficient range of motion (ROM) to articulate the joint to all desired positions. The *Mikrolar* robot noted in **Table 2** had the highest stiffness of any robot tested, which is not surprising, because high stiffness ratings is a notable benefit of hexapod robots [20]. However, hexapod robots have ROM limitations which limit the types of joints that can be tested. Other important attributes to consider include payload capacity, cost, and the ability to command the robots’ pose within the appropriate control system.

#### 4.2.2 Bone Stiffness

The AP stiffness of the tibia and femur shown in **Table 3** were within an order of magnitude of the average stiffness of all robots tested except for the parallel Mikrolar robots. Further, equation (9) shows the nonlinear relationship between bone length (*l*) and stiffness. This nonlinear relationship means that bone stiffness is highly coupled to length, and will decrease exponentially with bone length, the results of which can be seen in the System Load Error plots in **Figure 7**. The wide range of bone stiffness perpendicular to the long axis of the bone indicate that bone deformation in superposition testing is a significant consideration, in addition to robot deformation when there is an AP load applied.

#### 4.2.3 Joint Stiffness

The anterior stiffness of the knee joint itself is more difficult to quantify because it is a function of flexion angle and AP displacement of the tibia relative to the femur. Given the nonlinear force displacement curves, best described by a 3^rd^ order polynomial for knee laxity (**Figure 6A**), joint stiffness will also increase exponentially with displacement. However, the stiffness – displacement relationship appears more linear when plotting stiffness vs force (**Figure 6B**). **Figure 6B** demonstrates that the intact knee joint anterior stiffness range is generally between 10 to 70 N/mm when applying loads less than 100 N.

The SSM (31) and **Figure 7** both indicate that increasing the ratio of joint stiffness to fixture stiffness (such as the robot or bone stiffness) increases compliance induced superposition errors. Researchers cannot select joint stiffness, but robot stiffness and bone length can be controlled. When higher stiffness serial arm robots are being used, such as the Kuka KR300 in **Table 2**, robot stiffness can be an order of magnitude larger than AP joint stiffness at load ranges less than 100 N. The tibia and femur AP stiffness in **Table 3** is on the same order of magnitude as knee joint AP stiffness. In fact, both the tibia and femur can have similar stiffness ranges compared to the maximum knee joint stiffness (∼70 N/mm in **Figure 6**) with lengths greater than 21 cm or 25 cm respectively. This underscores the importance of minimizing bone length to minimize superposition computed System Load Error.

### 4.3 System Load Error

**Figure 7** provides a convenient estimate of error when computing ACL tension in previous biomechanical studies that do not compensate for fixture compliance. Depending on robot stiffness and bone length, errors can increase beyond 50%, which could obscure understanding of a ligament’s contribution to joint function. It is therefore very important that the appropriate robot and potting technique be used to minimize these System Load Errors.

Given the many assumptions needed to estimate that error, particular attention should be given to the relative relationship between system stiffness and error. As fixture stiffness decreases or joint stiffness increases, System Load Error increases, which agrees with SSM predictions. **Figure 8** further demonstrates this relationship by showcasing System Load Errors for specific robots, bone lengths, and joint angles.

**Figure 8:**
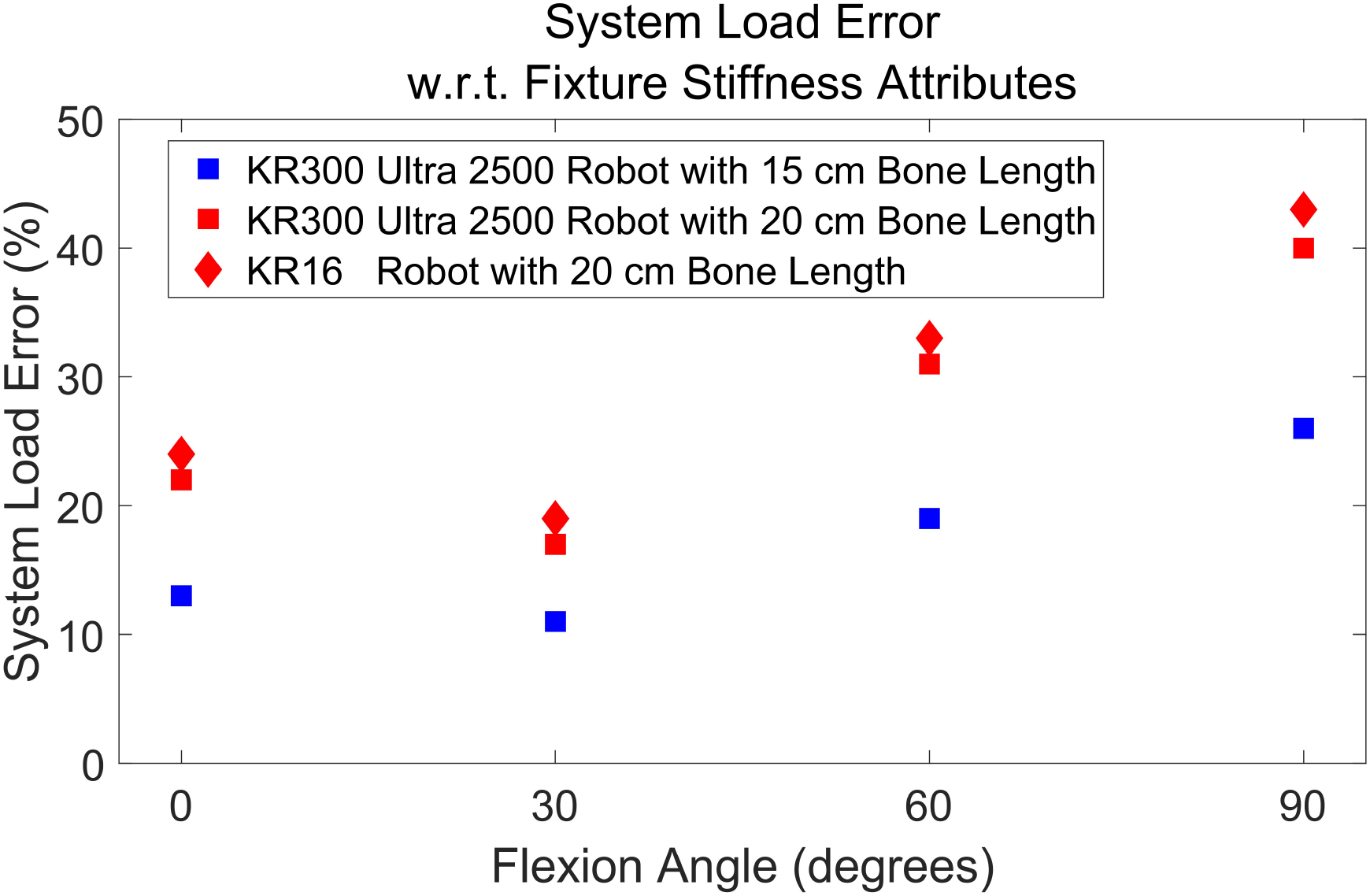
System Load Error for 2 common serial arm robots and bone lengths. Increasing bone length, decreasing robot stiffness, and varying joint stiffness by varying flexion angle all produce greater error.

Similar trends in System Load Error would likely emerge if other ligaments were analyzed. The magnitude of these errors will be dependent on the stiffness of the joint and ligament being tested. Further, the direction of loading during superposition also matters. If joint loading includes joint compression, then the joint stiffness may be much higher. For example, to understand the knee lateral collateral ligament’s role in joint stabilization, Varus-Valgus superposition might be performed. A Varus rotation will distract the lateral side of the joint while compressing the medial side, leading to coupling of this rotation with axial compression/distraction. While axial joint stiffness will not be as high as the bone’s axial stiffness (at least 1,390 ± 200 N/mm [18]) due to the soft tissue such as the meniscus and cartilage, the overall effect on System Load Error will likely increase in this DOF compared to AP.

There were three main limitations to the study methodology when quantifying system compliance and error. First, only the knee joint AP load contribution for the ACL was examined. This condition was selected to demonstrate the proposed framework for evaluating uncertainties reported in situ load contribution because it is one of the most studied topics using superposition testing [2-4, 6, 8, 10-13, 16, 17]. Second, while robot stiffness was characterized, it was difficult to quantify because it is a function of robot pose [20, 21]. When possible, standardized robotic home positions used for calibration purposes were used for all robots in this study, but in future testing it might be more correct to characterize robot stiffness at poses more similar to those typically observed in biomechanics testing. Third, in the calculation of System Load Error approximations of ACL contribution and ACL deficient knee stiffness were used because there were no known data of better quality. ACL contribution values were obtained from Spokane et al. and used to calculate *F*_*JSM*_ in (20), but these values are subject to the System Load Error of the test system used in that publication. An ACL intact knee model was used to model an ACL deficient knee in the calculation of System Load Error. This was done because the <u>Open Knee project</u> knee laxity data was available in the literature and was of high enough fidelity to fit 3^rd^ order polynomials to the data with good agreement [19]. There was no known ACL deficient knee laxity data of similar fidelity available in the literature. It is expected that the magnitude of the error calculations might differ if an ACL deficient model was utilized. However, the ACL intact knee model will still provide insight into the relative relationship between system stiffness and error, and that the magnitude of errors will be similar to a deficient knee.

Future work includes characterizing the stiffness of other robots, bones, and biological joints in multiple DOF. Characterizing the various stiffness may help to determine which robots might work best with which biological joints. Finally, an alternative approach to superposition testing is being evaluated which controls joint displacement directly (*ΔX*_*J*_) rather than system displacement (*ΔX*_*S*_) to perform the superposition testing shown in the SSM Deficient Loading Case in **Figure 3**. This approach enforces the principle of superposition on joint displacement in real-time by compensating for system compliance using 6-DOF position sensors [22].

## 5. CONCLUSION

To **explain** the mechanistic influence of system compliance on systemic positioning error in superposition testing, a system stiffness model was created. The model assigned a stiffness to every component of a testing system and demonstrated why ligament tension and stiffness have been historically underestimated when there is unaccounted compliance in a test system. To **quantify** the effect that positioning error has had in the biomechanics literature, stiffness of many components of a typical test system were measured. The stiffness of robots and bones was often in the same order of magnitude as the joint being tested. To **provide a framework** by which to evaluate uncertainty in published in situ force measurements based on superposition testing, System Load Error was calculated as a function of various robot, bone, and joint stiffness. The System Load Error demonstrates that the underestimation of ligament tension decreases with increasing system robot and bone stiffness, but increases with increasing joint stiffness. Depending on the robot setup and bone mounting protocol used System Load Errors may be substantial, as demonstrated by values of 50% or higher when calculated using published ACL forces.

It is recommended to minimize bone length and maximize robot stiffness when performing superposition testing in order to improve accuracy and reduce uncertainty of computed ligament tensions. Direct control of joint displacement, rather than system displacement, could also be a great alternative as this solution would not require sacrificing important robot attributes, like ROM. The ratio of joint stiffness to fixture stiffness (including robot or bone stiffness) can help determine the appropriate fixture stiffness for a given biological joint, loading direction and loading magnitude. In this study the System Load Error plots (**Figure 7)**, and the underlying methodology to compute System Load Error, provide insight into the prevalence of systemic error when conducting superposition methodology in various test systems. Future publications should report fixture stiffness when applicable, so that System Load Error might be evaluated.

Computer modelers, medical device designers, and clinicians rely on superposition testing to calculate the contribution of various ligaments to joint stability. More accurate estimates of ligament tension and stiffness will enable researchers and device manufacturers to better design and refine medical devices, and surgical techniques, in the future.

## ACKNOWLEDGMENT

The authors would like to acknowledge Lesley R. Arant and Joshua D. Roth of the University of Wisconsin-Madison for their technical support. In addition Mike Chaffee from Kuka Robotics was instrumental in taking stiffness measurements on the KR300 Ultra 2500.

## NOMENCLATURE

*F*_1_: Force 1: The force the system exerts on the intact joint in the JSM.
*F*_2_: Force 2: The force the system exerts on the deficient joint in the JSM.
*F*_A_: Force A: The calculated force held by Structure A.
*F*_*B*_: Force A: The calculated force held by Structure B.
*ΔX*: The system displacement in the JSM. This model makes no distinction between system and joint displacement and assumes they are the same.
*K*_*A*_: Structure A stiffness: The stiffness of structure A, as calculated by JSM. Because the JSM doesn’t distinguish between system and joint displacement, this stiffness a function of the system fixture and joint stiffness.
*K*_*B*_: Structure B stiffness: The stiffness of structure B, as calculated by the JSM. Because the JSM doesn’t distinguish between system and joint displacement, this stiffness a function of the system fixture and joint stiffness.
*ΔX*_1_: The system displacement of the intact joint in the JSM.
*ΔX*_2_: The system displacement of the deficient joint in the JSM.
*ΔX*_*J*_: Displacement of the joint from the unloaded position in the SSM.
*ΔX*_*FB*_: Displacement of the fixture base from the unloaded position in the SSM. *ΔX*_*FT*_ Displacement of the fixture tool from the unloaded position in the SSM. *ΔX*_*S*_ Displacement of the system from the unloaded position in the SSM.
*ΔX*_*S_*0_: Displacement of the system with an intact joint from the unloaded position in the SSM.
*ΔX*_*S_JSM*_: Displacement of the system with a deficient joint from the unloaded position when system displacement is controlled, as predicted by the SSM.
*ΔX*_*S_JSM*_: Displacement of the system with a deficient joint from the unloaded position when joint displacement is controlled, as predicted by the SSM
*EI*: Flexural Rigidity: The resistance of a structure (in this case bone) to bending.
*P*: Load applied to the bone.
*l*: Bone length
*CC*: Compliance Compensation Metric: the difference in joint displacement between the intact and deficient cases. It quantifies system positioning error.
*SLE (%)*: System Load Error: the calculated load error in structure A, as a percentage, if system compliance is not controlled for.
*K*_*JA*_: Structure A joint stiffness: The stiffness of Structure A, as calculated by the SSM. Because the SSM distinguishes between system and joint displacement, this stiffness the actual stiffness of Structure A.
*K*_*JB*_: Structure B joint stiffness: The stiffness of Structure B, as calculated by the SSM. Because the SSM distinguishes between system and joint displacement, this stiffness the actual stiffness of Structure B.
*K*_*F*_: Fixture stiffness: The stiffness of the tool and base fixtures combined, as calculated by the SSM.
*K*_*J*_: Joint stiffness: The stiffness of all structures in the joint, as calculated by the SSM.

## APPENDIX 1: SSM Derivation of Force and Displacement Distribution

An intuitive explanation for the distribution of displacement and force was given in Section 3.1, namely that [1] *F*_*JSM*_ < *F*_0_, [2] *ΔX*_*F _JSM*_ < *ΔX*_*F _*0_, [3] *ΔX*_*FT_JSM*_ < *ΔX*_*FT_*0_, [4] *ΔX*_*J_JSM*_ > *ΔX*_*J_*0_. Here these inequalities will be explicitly proved by describing the SSM using the equations for springs in parallel or series. Springs in series have the same force throughout, while springs in parallel must have their forces added. Therefore, F0 in the Intact Loaded Case of Figure 3 can be described by the SSM as:

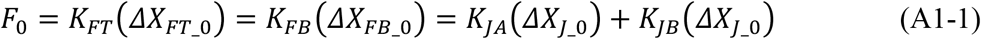

Finding the equivalent spring constant for the springs representing the fixture stiffness, which is in series, and the springs representing ligament stiffness, which are in parallel, (A1-1) can be written as:

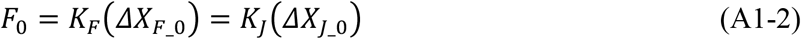

Where,

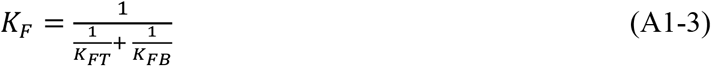

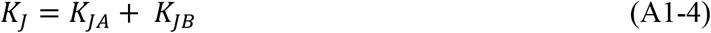

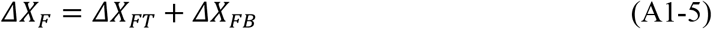

Similar to (A1-2), *F*_*JSM*_ from the **Figure 3** JSM deficient loaded case can be written as:

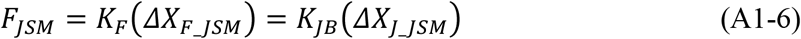

Writing (A1-2) as a ratio leaves us with:

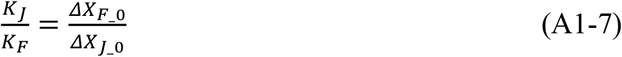

Similarly, writing (A16) as a ratio leaves us with:

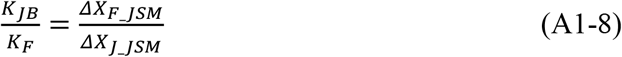

Substituting (A1-4) into (A1-8),

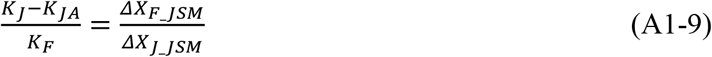

Given (A1-7) & (A1-9),

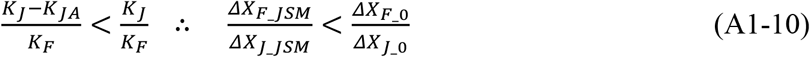

Substituting (7) & (A1-5) into (A1-10),

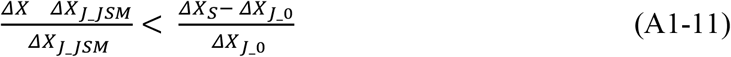

Rearranging this equation leads to the conclusion:

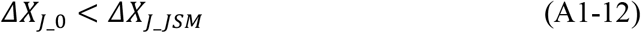

Given (7), (A1-5), & (A1-12),

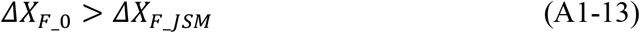

Substituting (A1-2) & (A1-6) into (A1-13),

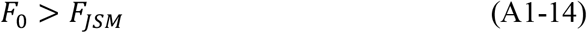

## APPENDIX 2: SSM Derivation of Error in Ligament Stiffness

The derivation of error in superposition computed ligament stiffness is more complex than the tension error derived in the results section. At the center of the stiffness error derivation is the concept that the Joint Stiffness Model assumes *ΔX* = *ΔX*_*J*_ *= ΔX*_*S*_. However, according to the SSM, because the JSM only measures actuator displacement, and fixture components are not infinitely stiff, only the assumption that *ΔX* = *ΔX*_*S*_ is true. Therefore using the terms from equations 4-6 and **Figure 3**, *ΔX*_1_ *= ΔX*_*S_*0_ and *ΔX*_2_ *= ΔX*_*S_JSM*_. The stiffness terms from the JSM equations 4-6 cannot be brought into alignment with the SSM stiffness terms because *K*_*A*_ and *K* in the JSM are a function of *ΔX*_*S*_ and therefore depend on both the stiffness of the joint structures and fixture stiffness *K*_*F*_. Therefore *K*_*A*_ ≠ *K*_*JA*_ and *K*_*B*_ ≠ *K*_*JB*_. Both models do agree that *ΔX*_*S_*0_ *= ΔX*_*S_JSM*_.

Both the intact and JSM cases of the SSM model control system displacement *ΔX*_*S*_. Similarly, in the JSM model, only system displacement is controlled. This implies that the system as a whole will be brought to the same displacement in both models. Therefore, it can further be assumed that *F*_0_ *= F*_*JSM*_ (4, 22) and *F*_2_ *= F*_*JSM*_ (5, 23).

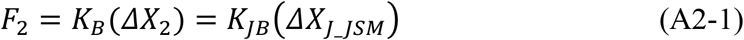

Using the definitions above (A2-1) may be rewritten as:

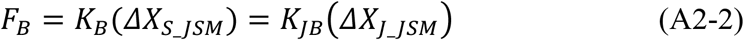

Substituting for *ΔX*_*J_JSM*_ using (7) from the SSM gives:

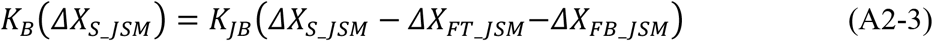

Rewriting,

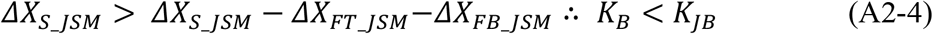

Because the JSM doesn’t distinguish between system and joint displacement, the system deflection in the JSM is larger than the joint displacement given by the SSM. To maintain the equality in (A2-3), the stiffness of Structure B as given by the JSM method must be smaller than what the SSM model predicts.

Given that *F*_0_ *= F*_*JSM*_ and *F*_2_ *= F*_*JSM*_, equations (6) and (24) may be written as:

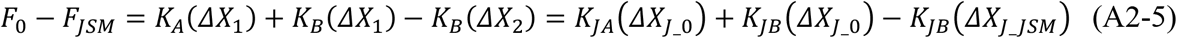

Given that *ΔX*_1_ *= ΔX*_2_ = *ΔX*_*S*0_ this simplifies to:

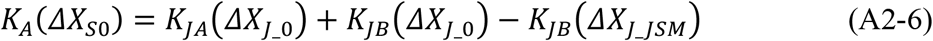

Further simplifying,

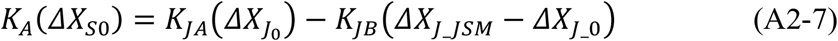

The Compliance Compensation Metric from (10) may then be substituted in:

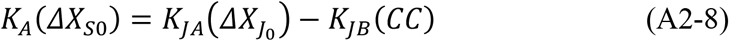

Rewriting (A2-8) by multiplying the K_JB_ term by 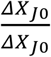 gets:

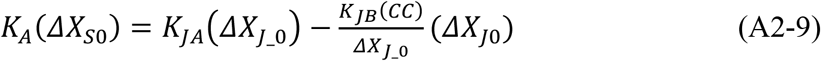

Rearranging,

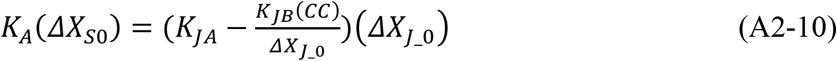

Using (7) & (A1-5),

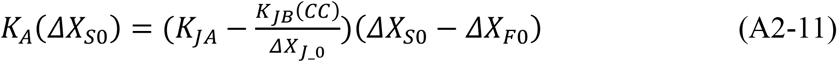

Using the same logic used in (A2-4),

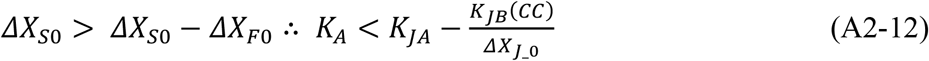

Because *CC* is always positive (A1-12), and the stiffness term *K*_*JB*_, and joint displacement term *ΔX*_*J_*0_ are also always positive, the joint stiffness (*K*_*JA*_) in (A2-12) is subtracted by a positive number. To maintain the equality in (A2-11), the stiffness of Structure A as given by the JSM method must be smaller than what the SSM model predicts Structure A joint stiffness with an additional term subtracted from it.

## Notes

### Competing Interest Statement

Callan Gillespie, Tara Nagle, and Robb Colbrunn received royalties from the sale of simVITRO robotic systems within the last 36 months.

